# A2A-positive neurons in the nucleus accumbens core regulate effort exertion

**DOI:** 10.1101/2024.12.14.628510

**Authors:** Charles T. Shoemaker, Alexander D. Friedman, Bryan Lu, Jiwon Kim, Shaolin Ruan, Henry H. Yin

## Abstract

Previous work has implicated the nucleus accumbens (NAc) in the regulation of effort, defined as the amount of work an animal is willing to perform for a given reward, but little is known about the specific contributions of neuronal populations within the NAc to effort regulation. In this study, we examined the contributions of direct pathway and indirect pathway neurons in the NAc core using an operant effort regulation task, in which the effort requirement is the number of lever presses needed for earning a food reward. Using optogenetics, we manipulated the activity of direct pathway spiny projection neurons (dSPNs, D1+) and indirect pathway SPNs (iSPNs, A2A+). Activating dSPNs reduced lever pressing regardless of the effort requirement, as it elicited gnawing, a competing consummatory behavior. On the other hand, activating iSPNs in the NAc core (but not in the shell) reduced lever pressing in an effort-dependent manner: stimulation-inducted reduction in performance was greater at higher press-to-reward ratio requirements. In contrast, optogenetically inhibiting NAc core iSPN output resulted in increased levels of effort exertion. Our results show that the indirect pathway output from the NAc core can bidirectionally regulate effort exertion.

## Introduction

The nucleus accumbens (NAc) is believed to play a fundamental role in regulating motivation. As a major component of the limbic basal ganglia circuit, it has long been thought to function as an interface between the neural substrates of motor control and limbic structures governing affect and homeostasis (Mogenson et al., 1980; Kelley, 2004; Yin et al., 2008; Nicola, 2010; Vancraeyenest et al., 2020).

Although the NAc is not necessary for learning action-outcome contingencies, it plays a major role in regulating instrumental performance (Corbit et al., 2001). Previous studies have established that the NAc is essential for the regulation of effort exertion (Aberman and Salamone, 1999; Kelley, 2004; Ghods-Sharifi and Floresco, 2010). Effort can be distinguished from motor output per se. It is defined as the amount of work an animal is willing to perform for a given amount of reward. Instrumental or operant conditioning provides a convenient way to quantify effort by measuring the number of lever presses the animal is willing to perform for a given amount of reward. For example, when a fixed number of presses (fixed ratio or FR) is required to earn a piece of food reward, the ratio represents the effort requirement.

Previous work has suggested that dopamine (DA) depletion in the NAc does not disrupt food consumption per se, but drastically reduces the number of lever presses for food rewards at high ratio requirements (Sokolowski and Salamone, 1998; Aberman and Salamone, 1999; Salamone et al., 2001). Depleted rats were still able to consume just as much home chow as normal rats, but they were much less willing to press when the ratio is high. Furthermore, rats with DA depletion in the NAc core show a reduced preference for effortful but large-reward actions (Mai et al., 2012). Such results suggest that mesolimbic DA may regulate effort exertion. It has also been shown that DA signaling in the NAc often reflects effort-related variables (Day et al., 2010; Ostlund et al., 2011).

Like the rest of the striatum, the NAc contains two major populations of spiny projection neurons (SPNs), which are typically classified as either direct (dSPNs) or indirect pathway SPNs (iSPNs). The direct pathway neurons express D1-like DA receptors, whereas the indirect pathway neurons co-express A2A adenosine and D2 DA receptors (Gerfen et al., 1990; Fink et al., 1992; Lu et al., 1997). Pharmacological studies have implicated the indirect pathway in regulating effort exertion. A2A agonists, which presumably increase the activity of iSPNs by activating G_s_-coupled A2A receptors and increasing neuronal excitability, disrupt performance on operant tasks with high effort requirements (Mingote et al., 2008). This finding suggests that increasing iSPN activity in the NAc core can reduce effort exertion without affecting general motor output. However, the role of the direct pathway in effort exertion is still unclear.

In the present study, we used optogenetics to test the contributions of the direct and indirect pathways by imposing distinct schedules of reinforcement and associated effort requirements in mice. FR schedules were used to manipulate the effort requirement without changing reward amount or delay. To specifically manipulate the activity of direct and indirect pathway neurons, optogenetic stimulation was done in Cre driver lines (D1-Cre for the direct pathway, and A2A-Cre for the indirect pathway). Our results suggest that activation of NAc core iSPNs reduces effort exertion. On the other hand, while stimulation of dSPNs also reduces lever pressing, it does so by eliciting competing consummatory behaviors like gnawing that interfere with lever pressing.

## Results

Initial operant experiments were conducted to determine whether the willingness of mice to exert effort would be impacted by optogenetically stimulating either core dSPNs or iSPNs. In order to selectively activate direct (D1+) or indirect (A2A+) pathway neurons in the NAc core, we used Cre-dependent channelrhodopsin (ChR2) and Cre driver lines that express Cre-recombinase under the promoters for the D1 or A2A receptors (Gerfen et al., 2013).

We first injected Cre-dependent ChR2 bilaterally in D1-Cre (n = 10, 3 males and 7 females) and A2A-Cre mice (n = 9, 4 males and 5 females), and implanted optic fibers targeting the NAc core. The control group consisted of wild-type (WT) mice that received the same injections and fiber implants (n = 5, 2 males and 3 females). Histological analysis of the mouse brains after the completion of all experiments confirmed that the fibers were implanted in the NAc core (**Figure 1A**).

**Figure 1.**
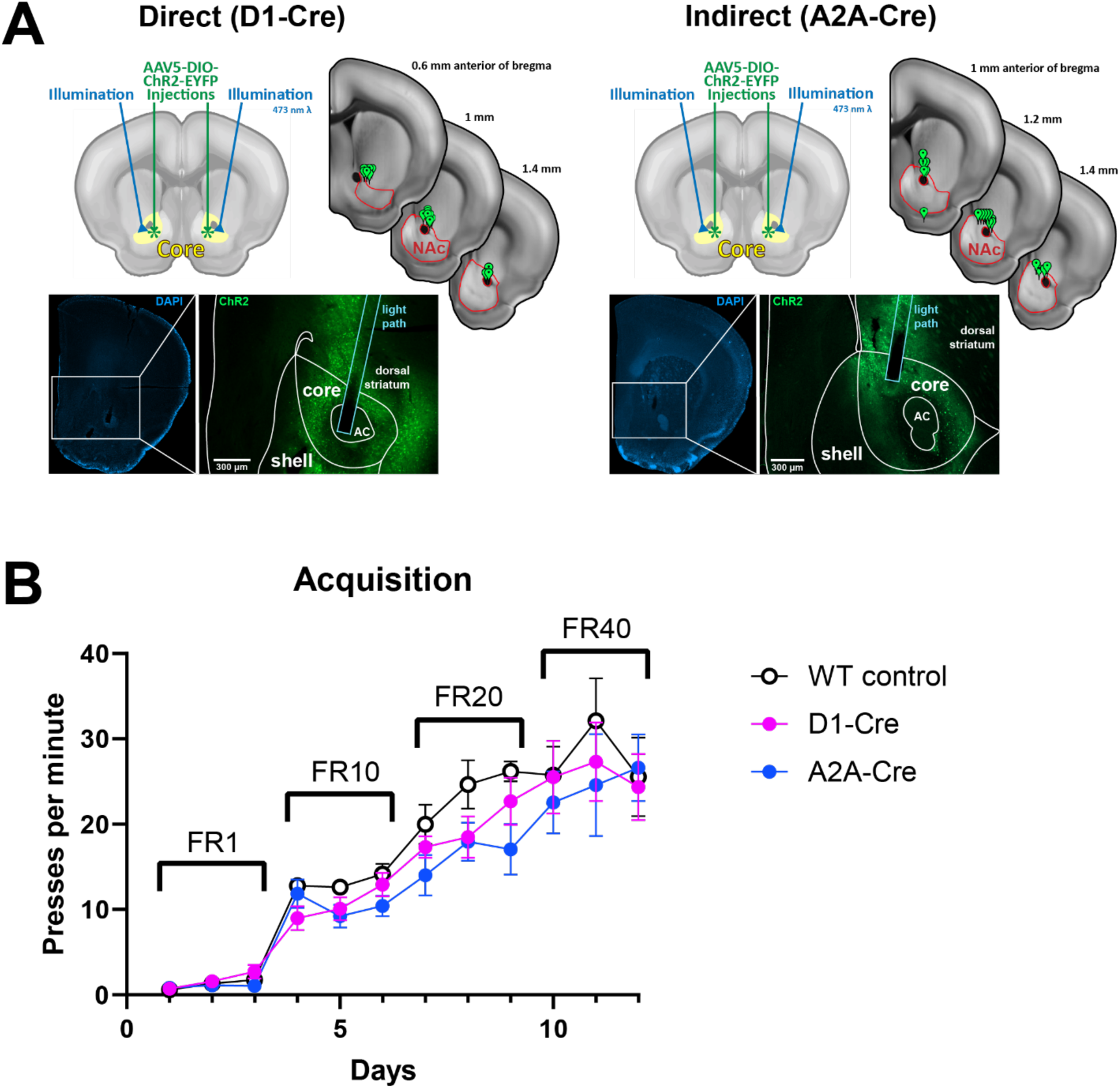
Virus expression, fiber placement, and operant training in mice expressing ChR2 in the NAc core. A) Expression of Cre-dependent ChR2 and placement of optic fibers in the NAc core. Left, fiber placement and representative histology for the direct pathway. Right, indirect pathway. B) Acquisition data from wild-type (WT) control, D1-Cre, and A2A-Cre mice. There were no group differences in acquisition. All three groups received the same virus injection in the NAc core. Error bars indicate SEM.

After recovering from surgery, all mice readily acquired lever pressing during FR training. Their rates of lever pressing during training sessions are shown in **Figure 1B**, and increased significantly for all three groups (main effect of session: F(11, 224) = 56.72, p < 0.0001).

During stimulation sessions, photo-stimulation immediately reduced lever pressing in both direct and indirect groups. From one epoch to the next, pressing reductions caused by stimulation were clearly observed after direct pathway activation (8 Hz stimulation data is shown in **Figure 2A**). Likewise, indirect pathway stimulation also reduced lever pressing (**Figure 2B**).

**Figure 2.**
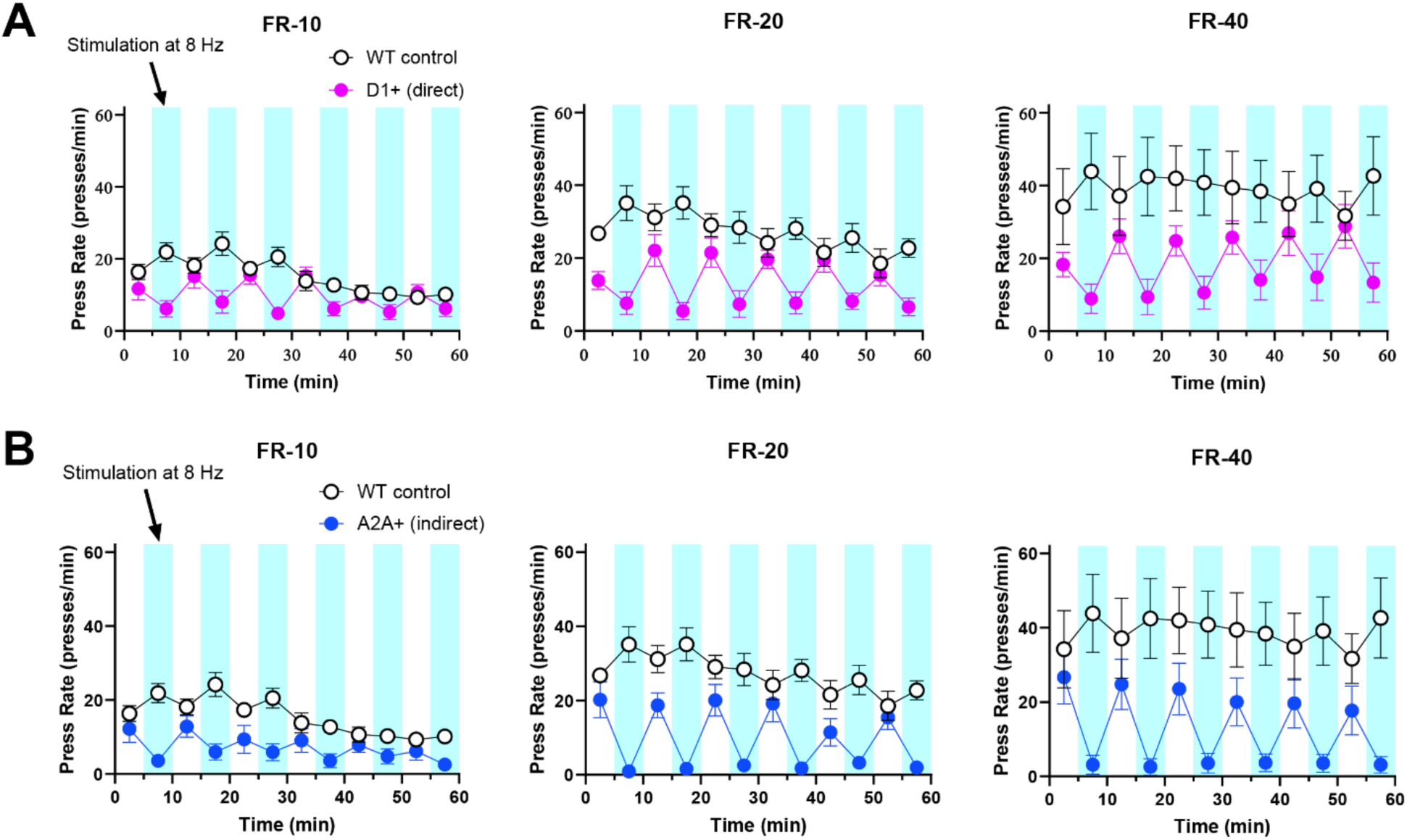
Time Course of Training and Optogenetic Testing for Operant Experiments. A) Activation of the D1+ (direct pathway) neurons in the NAC core reduced lever pressing. Blue bars indicate time of photo-stimulation (473 nm, ChR2, 8 Hz, 10 ms pulse width). B) Activation of the A2A+ (indirect pathway) neurons in the NAC core reduced lever pressing. Blue bars indicate time of photo-stimulation (473 nm, ChR2). Error bars indicate SEM.

**Figure 3** shows a summary of the change in lever pressing during stimulation sessions. For FR-10 sessions, a 2-way ANOVA revealed a significant interaction between genotype (WT, D1-Cre, and A2A-Cre) and frequency (5 and 8 Hz) (F(2, 21) = 5.660, p = 0.0108). Post hoc analysis showed no differences between groups in the 5 Hz condition, but the D1 (p = 0.0162) and A2A (p = 0.0135) groups showed a larger reduction in lever pressing than the WT group in the 8 Hz condition. For FR-20 sessions, the interaction was also significant (F(2, 21) = 5.027, p = 0.0164). Post hoc analysis showed that the reduction caused by 5 Hz stimulation in the A2A+ group was greater than in the control group (p = 0.0057). For the 8 Hz condition, both the D1+ (p = 0.0001) and the A2A+ groups (p < 0.0001) showed greater reductions in pressing compared to controls. In the FR-40 sessions, there was a significant main effect of genotype (F(2, 21) = 10.93, p = 0.0006). Post hoc analysis showed that reductions in pressing caused by 5 Hz were larger for the A2A group than for the control group, but this was not the case for the D1+ group (p = 0.0006). Reductions caused by 8 Hz in both the D1+ (p = 0.0026) and the A2A+ groups (p < 0.0001) were greater than in controls.

**Figure 3.**
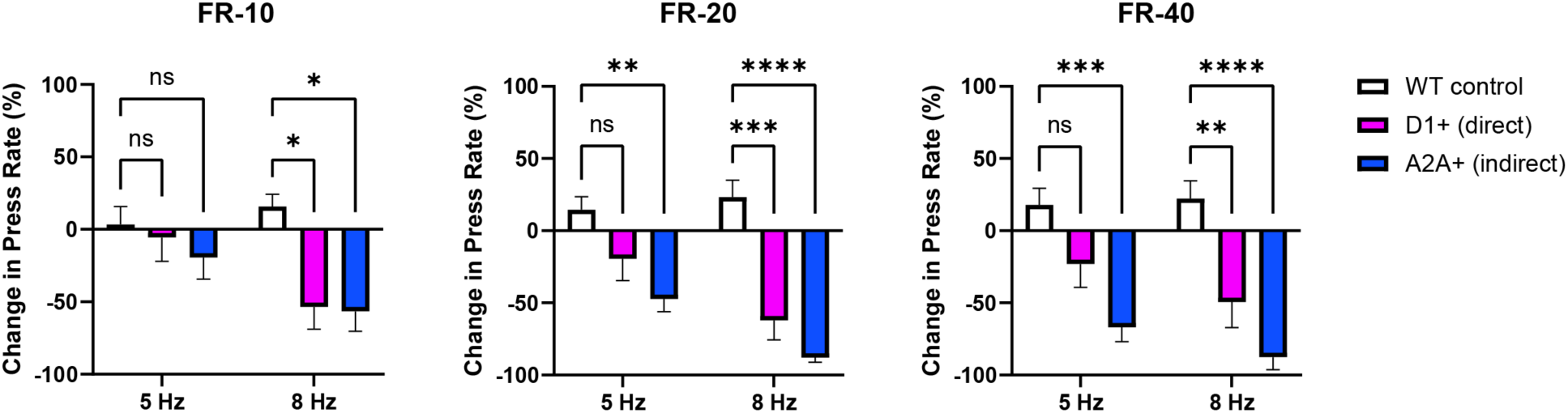
Summary of stimulation effects in D1+ and A2A+ neurons in the NAC core. ChR2 excitation of direct or indirect pathway neurons significantly reduced the rate of lever pressing. This effect is strongest at FR-40, suggesting that the % decrease in performance is greater when the effort requirement is high. * p < 0.05. ** p < 0.01. *** p < 0.001. **** p < 0.0001. Error bars indicate SEM.

To determine if the reduction in lever pressing observed depends on effort requirement, we plotted change in lever pressing as a function of the fixed ratio (the number of presses required per reward). As shown in Figure 4A, in the WT control group and the D1+ group, the reductions in pressing showed no significant relationship with the effort requirement (WT 5 Hz: R^2^ = 0.0262, p = 0.5644; WT 8 Hz: R^2^ = 0.0216, p = 0.6013; D1+ 5 Hz: R^2^ = 0.0168, p = 0.4943; D1+ 8 Hz: R^2^ = 0.0005, p = 0.9038). By contrast, in the A2A+ group, the percent change in lever pressing increases as the ratio is increased, as indicated by a significant linear relationship in the 5 Hz condition (R^2^ = 0.2338, p = 0.0106) and a marginally significant relationship in the 8 Hz condition (R^2^ = 0.1434, p = 0.0515).

**Figure 4.**
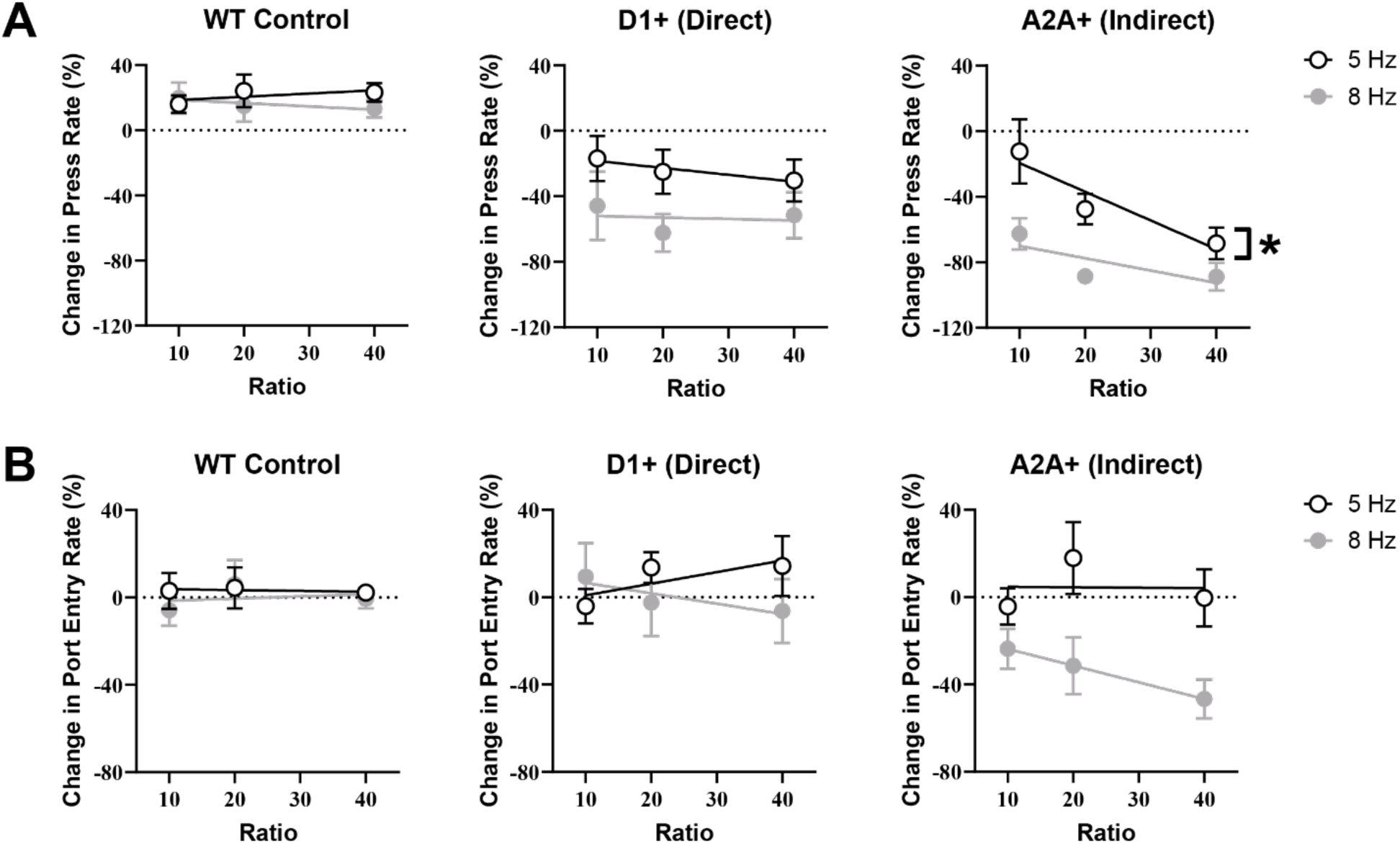
Change in press rate and reward port entry as a function of fixed ratio requirement. A) Effort-dependent reduction in lever pressing was found only after activation of A2A+ neurons in the NAC core: As ratio increases, % reduction in press rate also increases. Regression analysis shows that the reduction in pressing caused by 5 Hz stimulation significantly depends on increasing ratio requirement. This relationship was marginally significant in the 8 Hz condition (p = 0.0515) because stimulation produced a floor in press rate from FR-20 onwards. This effect was not observed in WT controls or D1+ mice. B) Note that ratio-dependent reduction in performance was not observed in the rate of reward port entry, which requires less effort. * p < 0.05. Error bars indicate SEM.

For comparison, we also plotted the change in the rate of food port entry (Figure 4B). When a pellet is dispensed, the mice must enter the food port to retrieve it. This behavior was measured using a beam break. Stable rates of port entry can be used as an indication that levels of motor coordination and appetite are consistent, and changes in this measure caused by changes in the ratio requirement are relatively minimal. Our results showed that changes in port entry rate did not depend on effort requirement (WT 5 Hz: R^2^ = 0.0010, p = 0.9117; WT 8 Hz: R^2^ = 0.0055, p = 0.7931; D1+ 5 Hz: R^2^ = 0.0452, p = 0.2593; D1+ 8 Hz: R^2^ = 0.0166, p = 0.4972; A2A+ 5 Hz: R^2^ = 0.00004, p = 0.9728; A2A+ 8 Hz: R^2^ = 0.0929, p = 0.1222). Thus, in all three groups, changes in the rate of food port entry showed no relationship with effort requirement (no significant effect of ratio). Regardless of the ratio schedule, mice maintained a similar motivation and capacity to check the food port for pellets.

These results suggest that only activation of the indirect pathway (A2A+) in the NAc core had a ratio-dependent effect, resulting in greater reduction in performance at higher ratios. But activation of the direct pathway reduced lever pressing consistently, regardless of effort requirement. In accord with this finding, we also observed significant gnawing behavior during direct pathway stimulation: when stimulated, D1+ mice often gnawed on the edges of the reward port. To quantify this behavior, we used DeepLabCut to track the position of the snout and determine when it was within a centimeter of the reward port edges(Mathis et al., 2018). We used time in this zone as a putative measure of gnawing. This analysis was done for a subset of D1+ (n = 4) and WT control (n = 5) mice whose operant behavior was recorded by a high-resolution video camera. From one epoch to the next, stimulation reversibly increased the amount of time D1+ mice spent gnawing on the reward port (8 Hz stimulation data is shown in Figure 5A). Figure 5B shows a summary of the change in gnawing due to stimulation. For the FR-20 and FR-40 schedules, 2-way ANOVAs revealed a significant main effect of genotype (FR-20: F(1, 7) = 19.63, p = 0.0030; FR-40: F(1, 7) = 19.63, p = 0.0030). Both 5 and 8 Hz stimulation caused D1+ mice to increase time spent gnawing versus WT controls (FR-20, 5 Hz: p = 0.0259; FR-20, 8 Hz: p = 0.0053; FR-40, 5 Hz: p = 0.0342; FR-40, 8 Hz: p = 0.0027). For the FR-10 schedule, there was no main effect of genotype (F(1, 7) = 3.289, p = 0.1126). However, 8 Hz stimulation caused D1+ mice to increase time spent gnawing versus WT controls, although the effect was marginally insignificant (p = 0.1064). This demonstrates that activation of the direct pathway reduced lever pressing by introducing a competing behavior: gnawing.

**Figure 5.**
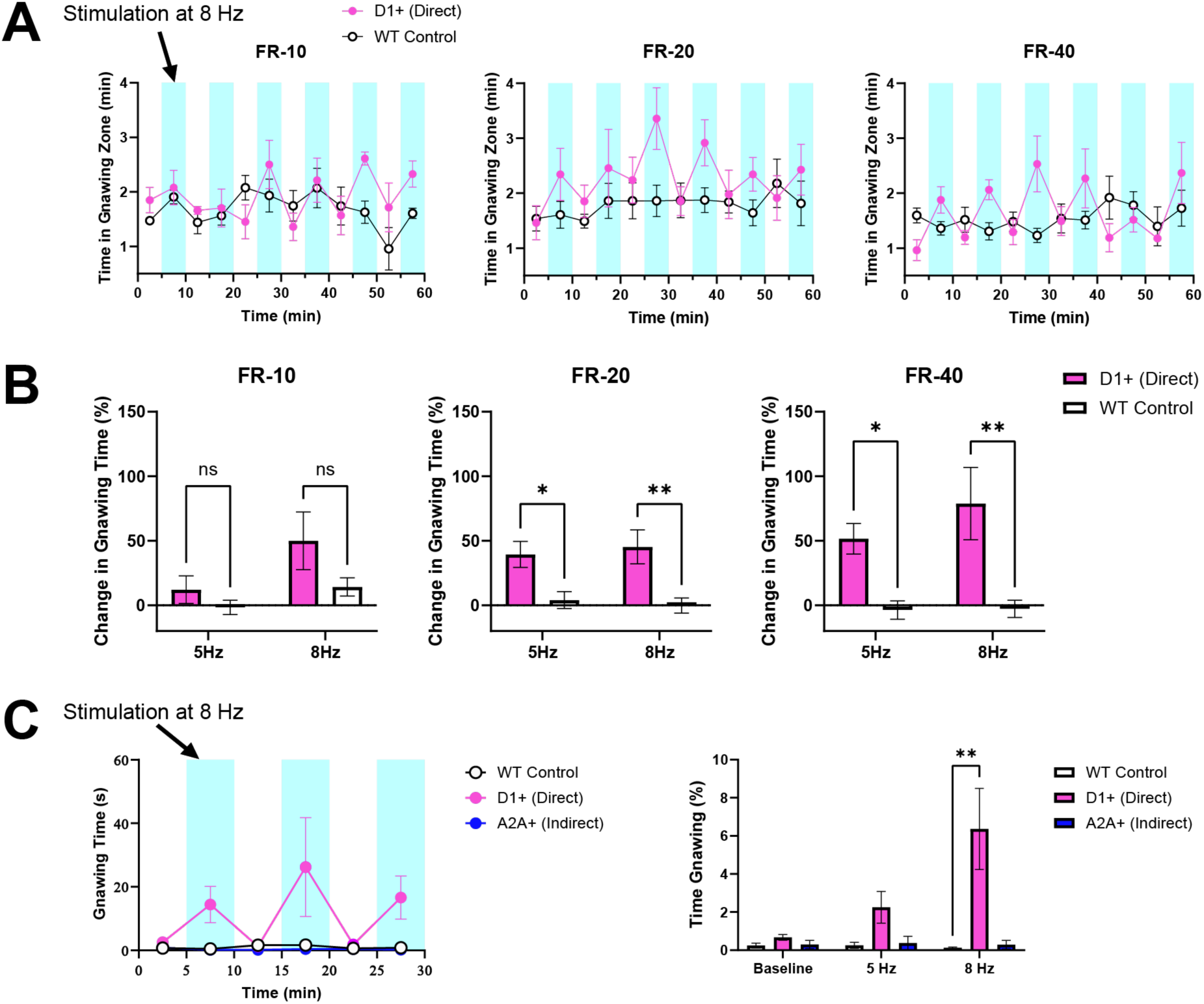
Activation of the direct pathway (D1+) causes gnawing, a consummatory behavior that competes with lever pressing behavior. A) Rapid and reversible increase in gnawing behavior elicited by optogenetic activation (8 Hz) of D1+ neurons in the NAC core during operant performance. B) Optogenetic stimulation in D1+ mice causes a significant increase in gnawing. C) Direct pathway activation also causes gnawing behavior in a separate chamber specifically designed to assay gnawing with high precision. *Left*: Rapid and reversible increase in gnawing behavior elicited by optogenetic activation (8 Hz) of D1+ neurons in the NAC core. In contrast, activating the indirect pathway (A2A+) had no effect on gnawing. *Right*: % time spent gnawing at different stimulation frequencies. There was a significant difference between D1+ and WT control mice in their percentage of time spent gnawing. * p < 0.05. ** p < 0.01. Error bars indicate SEM.

However, gnawing is best examined outside of the operant chamber, in an environment where an unobstructed view of the mouth can be obtained. Mice were also stimulated in a separate chamber, with a clear floor, where a high-resolution video camera could view the mouth from below. In agreement with our quantification of gnawing during operant behavior, activation of D1+ neurons produced gnawing in this context as well (**Supplementary Video 1)**. Activation of A2A+ neurons, on the other hand, did not produce gnawing (main effect of genotype: F(2, 14) = 5.306, p = 0.0193; Figure 5C). Thus, while activation of D1+ and A2A+ neurons in the NAc core can both reduce instrumental performance, activation of D1+ neurons elicits consummatory behaviors like gnawing that interfere with lever pressing, whereas activation of A2A+ neurons reduces lever pressing in an effort-dependent fashion.

To confirm that this effect is indeed specific to the core region, we also activated A2A+ neurons in the neighboring shell region of the NAc (Figure 6). This was done by injecting the same Cre-dependent ChR2 into the A2A-Cre (n = 8, 5 males and 3 females) and WT mice (n = 4, 4 females), but the virus and optic fibers were targeted to the shell region instead of the core. Unlike activation of A2A+ neurons in NAc core, activation of A2A+ neurons in the shell did not significantly affect behavioral measures: neither effort-dependent nor effort-independent reductions in lever pressing nor port entry were observed

**Figure 6.**
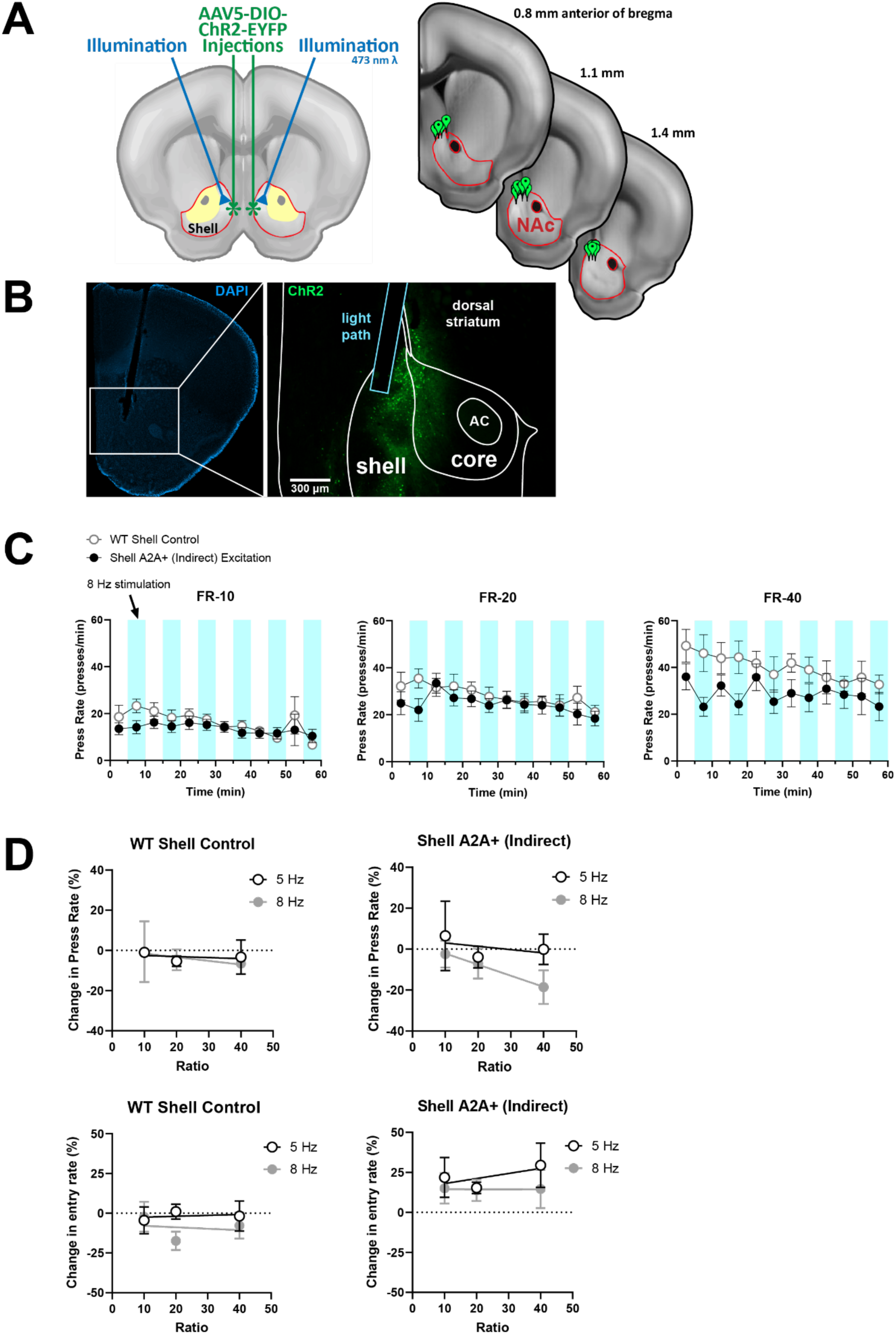
A) Fiber placement for optogenetic excitation of A2A+ neurons in the NAC shell. B) Representative slice showing virus expression in the NAc shell and optic fiber targeting the shell. C) Optogenetic excitation of A2A+ neurons in the NAC shell did not significantly reduce lever pressing. Data from NAC shell WT control group is shown for comparison. D) Changes in press rate (top) and reward port entry rate (bottom) as a function of the ratio requirement. WT control data on the left, A2A+ data on the right. Activation of A2A+ neurons in the NAC shell had no significant effect. Error bars indicate SEM.

If activation of the A2A+ neurons in the NAc core is responsible for suppressing effort exertion, inhibition of their activity could have the opposite effect of enhancing effort exertion. To test this prediction, we used the inhibitory opsin GtACR2 to reduce activity in A2A+ NAc core neurons (Mahn et al., 2018). The previously used A2A-Cre (n = 11, 5 males and 6 females) and WT mouse lines (n = 6, 2 males and 4 females) received the same optic fiber implants and GtACR2 injections. As shown in Figure 7, inhibition of the NAc core A2A+ neurons increased effort exertion as expected. There was no significant interaction between ratio and inhibition (F (2, 29) = 1.5, p = 0.24, no main effect of ratio F (2, 29) = 0.10, p = 0.90, and a main effect of inhibition (F (1, 15) = 5.12, p = 0.039).

**Figure 7.**
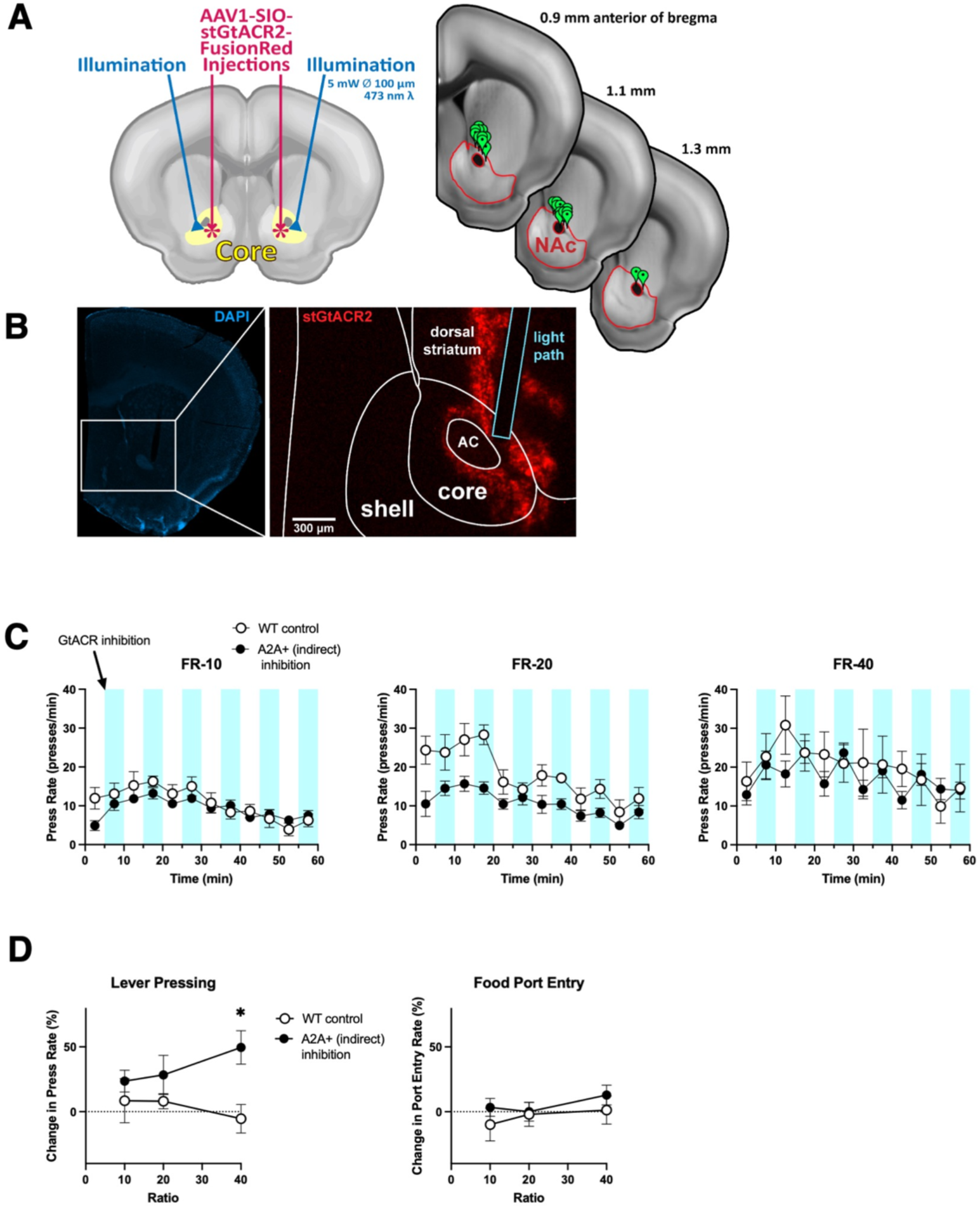
Optogenetic inhibition of A2A+ neurons in the NAC core increases lever pressing at higher effort requirements. A) Fiber placement for optogenetic inhibition with stGtACR2. B) Representative histology showing virus expression and optic fiber placement. C) Time course showing the effect of optogenetic inhibition of A2A+ neurons in the NAC core. D) At FR-40, inhibition of A2A+ neurons significantly increased lever pressing (* p = 0.0205; two-way ANOVA post hoc test). Error bars indicate SEM.

## Discussion

We showed that manipulation of neural activity in the direct and indirect pathways in the NAc core had distinct effects on motivated behavior. Lever pressing on FR schedules was significantly reduced in mice receiving either dSPN (D1+) or iSPN (A2A+) activation in the NAc core (Figure 3). However, there was a key difference between the effects of activating dSPN and iSPNs: dSPN activation reduced pressing to a similar extent regardless of the effort requirement, whereas iSPN activation resulted in a greater reduction in performance at higher ratios (Figure 4). Our experimental design allows us to dissociate general effects on performance due to changes in the motor system or appetite from more specific changes in effort exertion. Our results showed that the reduction in effort exertion during iSPN stimulation in the NAc core was not due to motor impairments or changes in appetite, as reward port entry or consumption was not affected in a ratio-dependent fashion.

Although the activation of both the direct pathway and the indirect pathway in the NAc core reduced lever pressing, the behavioral changes are distinct. Direct pathway stimulation elicited a dramatic increase in gnawing (Figure 5), and this effect accounts for reduced lever pressing, as gnawing behavior interferes with lever pressing. This finding suggests that direct pathway activation in the core can result in a switch to consummatory behaviors. Our findings also show that the contribution of the direct pathway is not the opposite of the indirect pathway. Although dSPN activation also reduced lever pressing, it did so independently of the effort requirement, producing similar reductions in performance regardless of the ratio requirement. The NAc has long been implicated in oral stereotypy (Delfs and Kelley, 1990). Previous work found that DA agonists, including D1 selective agonists, injected into the NAc could increase gnawing behavior (Tirelli and Witkin, 1995; Geter-Douglass et al., 1997). However, it is possible that the direct pathway still plays a role in effort regulation, but that the responsible dSPNs are found in different areas than the iSPNs. At any rate, dSPNs in the NAc core do not appear to play a role in regulating effort exertion as measured by lever pressing.

On the other hand, iSPN activation did not elicit gnawing behavior, but simply suppressed lever pressing, which was resumed as soon as stimulation stopped. Our results suggest that, in the NAc core, the dSPNs do not directly regulate effort exertion but mediate consummatory behavior. On the other hand, iSPNs play a critical role by suppressing performance when the effort required or the behavioral cost is too high. When the effort required is low, as in reward port entries or consumption, iSPN activation had limited impact on performance.

In addition, we found that the effect on effort exertion is restricted to iSPNs in the core. Stimulation of iSPNs in the neighboring shell region did not have a significant effect on effort exertion (Figure 6). Previous work has shown that DA depletion in the core, rather than the shell, was responsible for regulating performance (Sokolowski and Salamone, 1998). Thus, our results confirm these observations and suggest that the NAc shell is not a critical site for effort regulation.

A previous study showed that activation of either D1- or D2-SPNs in the NAc is sufficient to increase operant performance in rats (Soares-Cunha et al., 2016), whereas we found that indirect pathway activation reduces rather than increases effort exertion. However, it should be noted that our experimental designs and methods differed significantly from those in the study by Soares-Cunha and colleagues. In particular, we did not rely on the D2 promoter to express opsins. Because D2 receptors are expressed highly in interneurons such as the cholinergic interneurons, D2 may not be the best marker for iSPNs. In contrast, the expression of A2A receptors is limited to D2+/A2A+ SPNs, so we used Cre-dependent vectors in A2A-Cre mice.

The indirect pathway originating in the NAc core projects to the ventral pallidum. Previous work showed that inhibition of iSPN terminals to the ventral pallidum is sufficient to enhance performance, in accord with our optogenetic inhibition results (Nowend et al., 2001). Whereas activation of these iSPNs in the NAc core reduces effort exertion, inhibition of iSPNs can have the opposite effect and increase lever pressing, especially at FR-40 (Figure 7). While the increase in performance is more modest compared to the dramatic decrease observation during optogenetic excitation, this could be a ceiling effect, as the mice are already exerting considerable effort at higher FR schedules. Moreover, during high effort tasks like FR-40, the iSPN output may already be so low that inhibition can only boost performance to a relatively moderate extent. However, electrophysiological measures will be needed to test this possibility.

Our findings are also in agreement with previous work in rats showing that pharmacological manipulations of iSPNs using A2A agonists could reduce effort exertion (Mingote et al., 2008). Since A2A agonists are expected to increase iSPN activity, the effect could be similar to that of the optogenetic excitation used in our experiments. On the other hand, A2A antagonists, which like optogenetic inhibition can reduce iSPN activity, are known to increase effort exertion, as measured by lever presses and highest ratio achieved on a progressive ratio task (Randall et al., 2012).

Compared to previous pharmacological manipulations, a major advantage of our study is the temporal and cell-population selectivity. We are therefore able to demonstrate rapid and reversible changes in lever pressing by manipulating iSPN activity. We are also able to demonstrate bidirectional regulation of effort exertion by showing the opposite effects of excitation and inhibition on the same behavioral measure. In addition, our finding that the NAc core, but not the neighboring shell region, is critical for effort regulation is also in accord with previous work implicating the NAc core but not NAc shell as part of a neural circuit for effort regulation (Ghods-Sharifi and Floresco, 2010).

Our results are also in agreement with previous functional magnetic resonance imaging (fMRI) studies in humans (Botvinick et al., 2009). The magnitude of NAc activation was found to vary with both reward outcome and the effort demanded to obtain individual rewards. For a fixed level of reward, the NAc was less strongly activated following high effort demand than following low demand. This effect is correlated with preceding activation in the dorsal anterior cingulate cortex, which is thought to contribute to effort regulation. While intriguing, fMRI studies cannot reveal the specific neuronal populations involved. Recent work has shown that the activation of anterior cingulate projections to the dorsomedial striatum can reduce effort exertion (Ulloa Severino et al., 2023). As the anterior cingulate also projects to the NAc core, it is possible that its projections to the iSPNs can moderate effort exertion as well. One possibility is that cortical representations of behavioral cost are used to activate the anterior cingulate-NAc core iSPN circuit and reduce effort exertion. This hypothesis remains to be tested.

## Methods

### Animals

Experiments were conducted using adult mice (n = 53, 3-8 months), housed individually or in same-sex groups of no more than four. Three genotypes were used: D1-Cre mice (Drd1a-Cre, GENSAT, MMRRC_030989-UCD, n = 10, 3 males, 7 females), A2A-Cre mice (Adora2a-cre, MMRC_036158-UCD, n = 28, 14 males, 14 females), and wild-type control animals (C57BL/6J, Jackson Labs, n = 15, 4 males, 11 females). Mice had free access to water, except during experimental sessions. While undergoing food restriction, mice were weighed daily and then provided with chow to maintain approximately 85% ad libitum body weights. All procedures were approved by Duke University’s Institutional Animal Care and Use Committee (protocol A162-22-09).

### Surgery

Before intracranial surgeries, general anesthesia was initiated via isoflurane inhalation, and mice were placed upon a stereotaxic surgical apparatus. Mice received intracranial injections of either a Cre-dependent ChR2 vector (2×10^13^ vg/mL, pAAV5-Ef1a-DIO- hChR2(E123T/T159C)-EYFP, Addgene_35509) or a Cre-dependent GtACR2 vector (4.2-21×10^11^ vg/mL, pAAV1-hSyn1-SIO-stGtACR2-FusionRed, Addgene_105677). To target the NAc core (n = 36), eight injections (4 sites per hemisphere, 50 nL each) were administered targeting the following coordinates: 1 mm AP, 1.3 mm ML, and 4 mm DV; 1 mm AP, 1.3 mm ML, and 4.2 mm DV; 1.4 mm AP, 1.3 mm ML, and 4 mm DV; and 1.4 mm AP, 1.3 mm ML, and 4.2 mm DV. To target the shell region (n = 5), the following coordinates were used: 1 mm AP, 0.4 mm ML, and 3.8 mm DV; 1 mm AP, 0.4 mm ML, and 4 mm DV; 1.4 mm AP, 0.4 mm ML, and 3.8 mm DV; and 1.4 mm AP, 0.4 mm ML, and 4 mm DV. Subsequently, optic fibers measuring 100 μm in diameter were inserted at an angle of 10° from vertical (core coordinates for fiber tips: 1.2 mm AP, 1.4 mm ML, and 3.8 mm DV; shell coordinates: 1.2 mm AP, 0.8 mm ML, and 3.7 mm DV).

### Lever Press Training

Several weeks after surgery, food restriction was initiated, and mice were trained for an hour per day in 18×22 cm operant chambers (Med Associates). Mice were first trained on a fixed ratio 1 (FR-1) schedule of reinforcement: a 14 mg food pellet (Bio-Serv) was dispensed after each press of the single accessible lever. Timestamps were recorded for each lever press and each break of an infrared beam for detecting food port entries. After initial FR-1 training, each mouse was trained on an FR-10 schedule of reinforcement; in other words, a pellet was dispensed after 10 presses. After three days of FR-10 training, two daily optogenetic sessions tested 5 Hz and 8 Hz stimulation frequencies. This course of training and testing was then repeated for FR-20 and FR-40 schedules of reinforcement.

### Optogenetic Experiments

Optogenetic experiments were conducted using microcontroller-driven (Arduino, SCR_017284) 473 nm DPSS lasers. For inhibition experiments (n = 17), constant illumination was delivered during six alternating 5-min epochs starting in 10-min intervals in hour-long sessions. For excitation experiments (n = 36), 10 ms pulses were delivered during the same 5-min epochs. 5 Hz and 8 Hz stimulation frequencies were tested during two daily sessions. This was repeated for each ratio/effort requirement. For inhibition experiments, the laser power was approximately 5 mW in the brain. For excitation experiments, the laser power was approximately 3-4 mW, but an effect was elicited in one A2A+ NAc core mouse with 1 mW.

### Gnawing Measure

Inside the operant chamber, gnawing was quantified using DeepLabCut and Matlab (MathWorks). A high resolution video camera (Blackfly S, Teledyne FLIR) recorded mice from above. DeepLabCut identified the corners of the reward port and tracked mice’s snouts over time. A custom Matlab script used the corner locations to create a zone around the bottom edge of the reward port (~ 10 mm diameter), as well as the bottom half of the left and right edges. When the snout was within the zone, the behavior was classified as gnawing.

Outside the operant chamber, gnawing was assayed in a 22×22 cm enclosure with a transparent floor. One to four loops of black polylactide filament (1.5 mm in diameter, extending roughly 4 cm), on which mice would be able to gnaw, were fixed at each corner of the enclosure. Optogenetic manipulations were performed in an identical manner to that previously described, except that these sessions lasted 30 min, and that two sessions testing 5 Hz and 8 Hz stimulation frequencies were administered on a single testing day. To quantify gnawing, video was recorded from a camera below the enclosure, and the video data was analyzed with DeepLabCut and Matlab. DeepLabCut tracked the position of the mouth, and Matlab identified the loop locations. Behavior was classified as chewing when the mouth was within 2.5 mm of the loop.

### Histology

Mice were transcardially perfused with 0.1 M phosphate buffered saline and then 4% paraformaldehyde. Brains were extracted and submerged in 4% paraformaldehyde for at least 4 days and then in 30% sucrose for at least 2 days. 60 μm coronal sectioning was then performed via cryostat, and sections were mounted with an aqueous fluoroshield DAPI mounting medium (Abcam, Ab104139) and imaged (ZEISS Axio Zoom.V16, SCR_016980).

**Figure 8.**
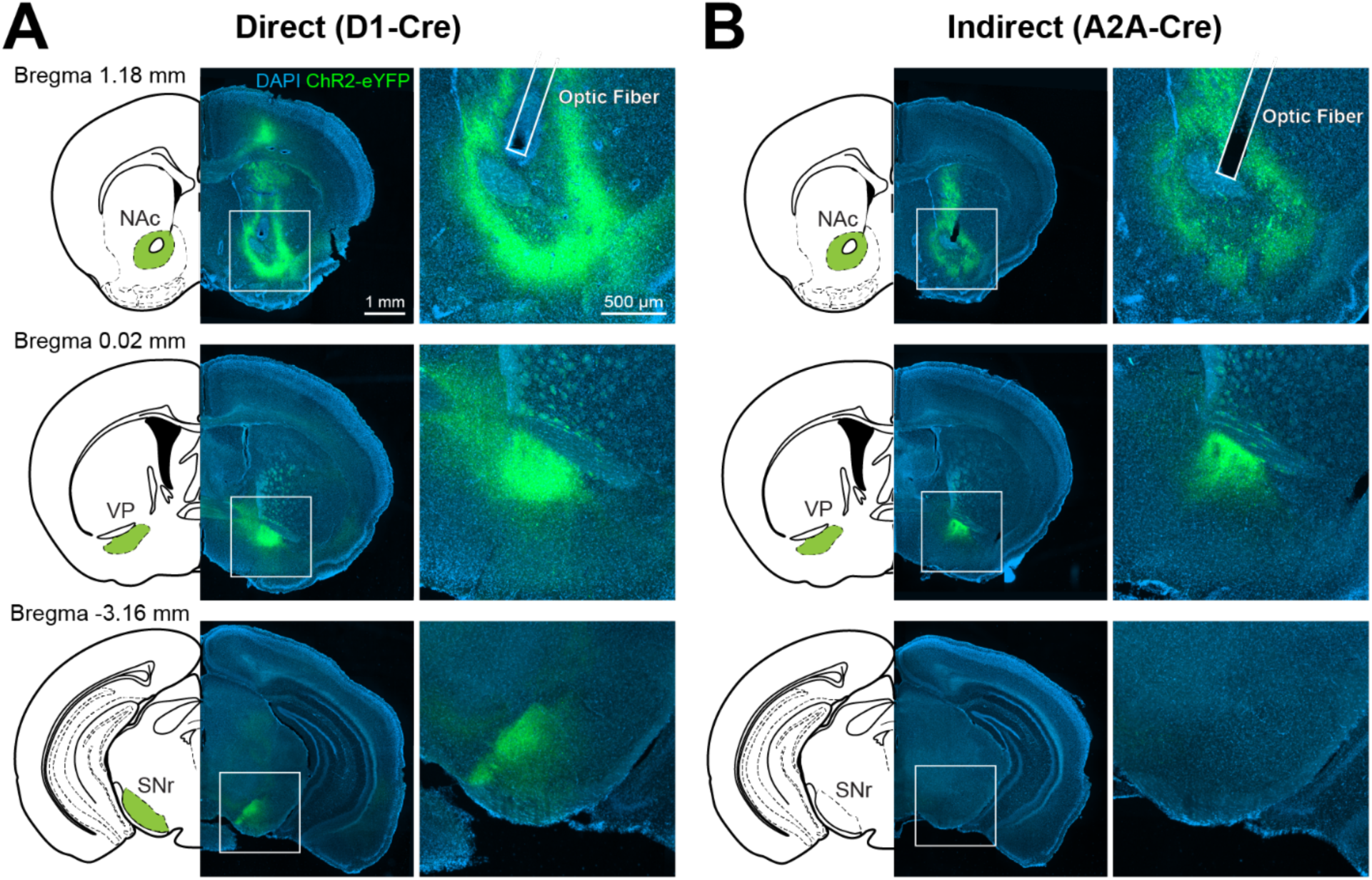
NAC core pathway-specific viral expression. A) Cre-dependent ChR2 expression in the direct pathway. D1+ neurons project to the ventral pallidum (VP) and substantia nigra pars reticulata (SNr). B) Cre-dependent ChR2 expression in the indirect pathway. A2A+ neurons project to the VP but not SNr. Same imaging parameters for D1+ and A2A+ neurons.

**Supplementary Video 1.** Gnawing behavior in a representative D1-Cre mouse expressing ChR2 in direct pathway neurons with 8 Hz stimulation in the NAc core.

